# Intranasal Administration of Functionalized Soot Particles Disrupts Olfactory Sensory Neuron Progenitor Cells in the Neuroepithelium

**DOI:** 10.1101/2020.08.19.256297

**Authors:** Jordan N. Norwood, Akshay P. Gharpure, Raju Kumal, Kevin L. Turner, Lauren Ferrer Pistone, Randy Vander Wal, Patrick J. Drew

## Abstract

Exposure to air pollution has been linked to the development of neurodegenerative diseases and anosmia, but the underlying mechanism is not known. Additionally, the loss of olfactory function often precedes the onset of neurodegenerative diseases. Chemical ablation of olfactory sensory neurons blocks the drainage of cerebrospinal fluid (CSF) through the cribriform plate and alters normal CSF production and/or circulation. Damage to this drainage pathway could contribute to the development of neurodegenerative diseases and could link olfactory sensory neuron health and neurodegeneration. Here, we investigated the impact of intranasal treatment of combustion products (laboratory-generated soots) and their oxygen functionalized derivatives on mouse olfactory sensory neurons, olfactory nerve cell progenitors, and the behavior of the mouse. We found that after a month of every-other-day intranasal treatment of soots, there was minimal effect on olfactory sensory neuron anatomy or exploratory behavior in the mouse. However, oxygen-functionalized soot caused a large decrease in globose basal cells, which are olfactory progenitor cells. These results suggest that exposure to air pollution damages the olfactory neuron progenitor cells, and could lead to decreases in the number of olfactory neurons, potentially disrupting CSF drainage.

## Introduction

Air pollution, particularly small combustion particles (<2.5 µm, PM_2.5_), is a large contributor to global mortality (Burnett et al., 2018). These small particles are produced by combustion in internal combustion engines, jet aircraft engines, and during cooking. Once generated, these particles can be oxidized over time (Rattanavaraha et al., 2011; Li et al., 2013; Pourkhesalian et al., 2015), generating surface functionalized oxygen groups which can increase their cellular toxicity (Li et al., 2009; Holder et al., 2012; Li et al., 2013).

In addition to the many other adverse health effects of air pollution, there is a strong epidemiological link between exposure to air pollution, particularly PM_2.5_, to the development of neurodegenerative diseases (Wang et al., 2017; Forman and Finch, 2018; Peters et al., 2019) and to mental disorders (Atanasova et al., 2008; Hummel et al., 2017; Buoli et al., 2018). Exposure to air pollution, particularly fine particulate matter (PM_2.5_), also leads to reduced sense of smell and anosmia (Ajmani et al., 2016a; Ajmani et al., 2016b) and can damage nasal tissue (Calderon-Garciduenas et al., 2003). Interestingly, anosmia and a decline of sense of smell precede the onset of neurodegenerative disorders (Doty, 1989; Wilson et al., 2009; Rahayel et al., 2012; Growdon et al., 2015; Ottaviano et al., 2016; Roberts et al., 2016; Murphy, 2019) and is also associated with depressive disorders (Croy et al., 2014; Kohli et al., 2016). Similar damage and sensory deficits have been implicated in COVID-19 pathology (Cooper et al., 2020). The observed associations between particulate exposure, decreased olfactory function, and development of neurodegenerative and mental disorders suggests that some of the observed degeneration might originate from the damage to olfactory sensory neurons (OSNs) in the nasal epithelium.

The movement of cerebrospinal fluid (CSF) is thought to remove waste from the brain (Iliff et al., 2012; Nedergaard, 2013), and disruption of normal CSF turnover and circulation has been hypothesized to lead to the development of neurodegenerative diseases (Albeck et al., 1998; Stoquart-ElSankari et al., 2007; Simon and Iliff, 2016; Benveniste et al., 2017). In addition to CSF drainage pathways through meningeal lymphatics and arachnoid granulations (Boulton et al., 1996; Aspelund et al., 2015; Louveau et al., 2016; Absinta et al., 2017; Ma et al., 2017; Ma et al., 2018), CSF is known to drain out of the cranial compartment via the cribriform plate, the perforated bone between the olfactory bulbs and nasal cavity (Bird et al., 2018), in both humans (de Leon et al., 2017) and animals (Bradbury et al., 1981; Mollanji et al., 2002; Nagra et al., 2006; de Leon et al., 2017; Norwood et al., 2019). The level of amyloid beta oligomers in this discharge is correlated with the severity of cognitive decline (Yoo et al., 2020). The fluid flows in-between olfactory neuron axons, and chemical ablation of OSNs blocks this normal outflow, leading to decreased CSF production and/or altered CSF circulation (Norwood et al., 2019). Thus, any damage to olfactory sensory neurons by air pollutants, in addition to impairing the sense of smell, might lead to disruption of normal CSF circulation which can then contribute to the development of neurodegenerative diseases.

Olfactory sensory neuron cell bodies are located in the nasal epithelium and send their axons to the olfactory bulb through the holes (foramina) in the cribriform plate. Because these neurons are exposed to the environment, they have a relatively short lifetime (several months (Gogos et al., 2000)), and are constantly replenished throughout the lifetime of the organism. Olfactory sensory neurons are generated from a population of nearby stem cells (Brann and Firestein, 2014; Liberia et al., 2019), and the ongoing neurogenesis of olfactory sensory neurons continues throughout the life of the animal. There are two classes of stem cells in the nasal epithelia that give rise directly and indirectly to OSNs, horizontal basal cells (HBCs) and globose basal cells (GBCs) (Child et al., 2018). GBCs generate olfactory sensory neurons, while HBCs are usually quiescent and are involved in regenerating the nasal epithelial in response to injury. The capacity for regeneration has limits and is reduced with aging or repeated insults (Child et al., 2018). Chronic nasal inflammation causes degeneration of olfactory neurons and their progenitor cells in both humans and animals (Chen et al., 2019; Hasegawa-Ishii et al., 2019). Insults that kill olfactory sensory neurons and their progenitor cells will lead to shrinkage and loss of the nasal CSF outflow pathways. Insults that kill either GBCs or HBCs will decrease the population of stem cells, potentially resulting in a decrease in the number of OSNs later in life.

To better understand the effects of air pollution on olfactory sensory neurons and their progenitor cells, we investigated the impact of intranasal treatment with surrogates for combustion generated ‘soots’ synthesized from carbon black precursors. Carbon black is primarily composed of elemental carbon, but like combustion-produced soot, it is formed by the partial combustion or thermal decomposition of hydrocarbons (Donnet, 1993). The morphology consists of primary particles that are partially merged or appear “fused” into aggregates (Fig. 1A). Such synthetic soots are also free of variable combustion-derived contaminants such as metals, ash, or condensed organics. We treated the mice with either non-functionalized soots, which resemble the combustion products immediately after their production, or functionalized soots that have been subject to oxygen functionalization, modifying their surface chemistry, mimicking the oxidation processes that would take place during atmospheric aging. We found that relative to vehicle controls, neither non-functionalized soots nor oxygen-functionalized soots had appreciable impact on olfactory sensory neurons. The effects of soot exposure on exploratory behavior was also minimal. However, oxygen-functionalized soots greatly decreased the levels of olfactory progenitor cells, suggesting that exposure to these particles can set up a long-term decrease in the number of OSNs. Such a decrease could lead to anosmia and decreased CSF movement.

**Figure 1:**
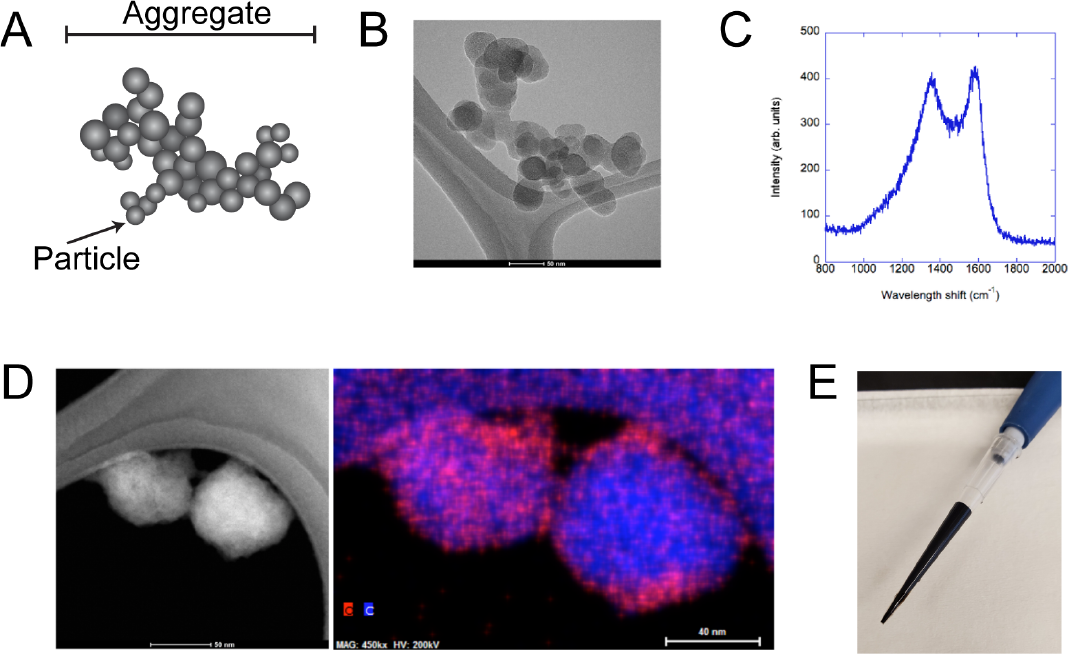
Soot characterization. A) Schematic showing the structure of a soot particle, which is an aggregate of smaller particles. B) TEM image of a soot aggregate supported by the lacy mesh of the TEM grid, illustrating the morphological structure of the particles. Pseudo-spherical primary particles are coalesced, forming a branched aggregate whose 2D projection is shown in the image. C) Raman spectrum of R-250 carbon black. The two peaks of similar intensity are indicative of unstructured (non-graphitic) carbon. D) Left, a high angle, annular dark field (HAADF) image of soot particles. Formed by scattered (rather than transmitted) intensity, the uniformity illustrates the lack of crystallinity and absence of heavy elements such as metals. Right, corresponding energy dispersive spectroscopy (EDS) map. The elemental map reveals the spatial distribution of carbon and oxygen, integrated through the particle. The higher intensity at the particle perimeters shows that the oxygen is at the particle surfaces. The grid lacy mesh appears as arched support). E) Photograph of suspended soot (1%) in water. The oxygen surface functionalization makes the particle hydrophilic, enabling stable dispersion in aqueous media.

**Figure 2:**
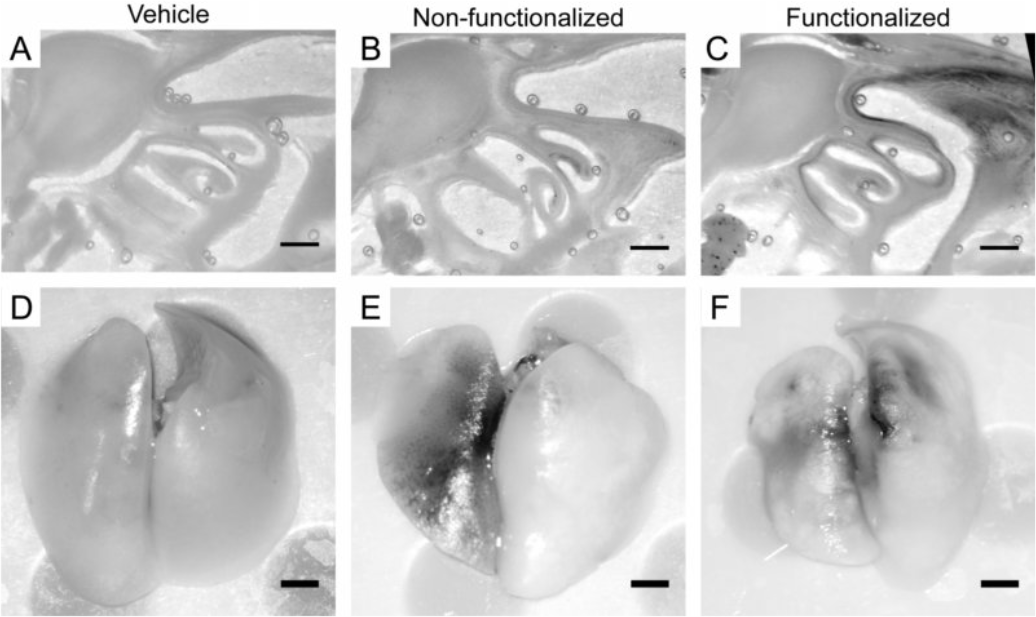
Uptake of soot in nasal epithelium and lungs. Images of sagittal sections of decalcified skull and nasal epithelium after one month of treatment with a vehicle (A), non-functionalized soot (B) and functionalized soot (C). Images of the lungs of vehicle-treated (D), non-functionalized (E) and functionalized soot-treated (F) mice. Only the functionalized soot shows appreciable accumulation in the nasal epithelium, though both soot types accumulate in the lungs. All scale bars 1 mm.

**Figure 3:**
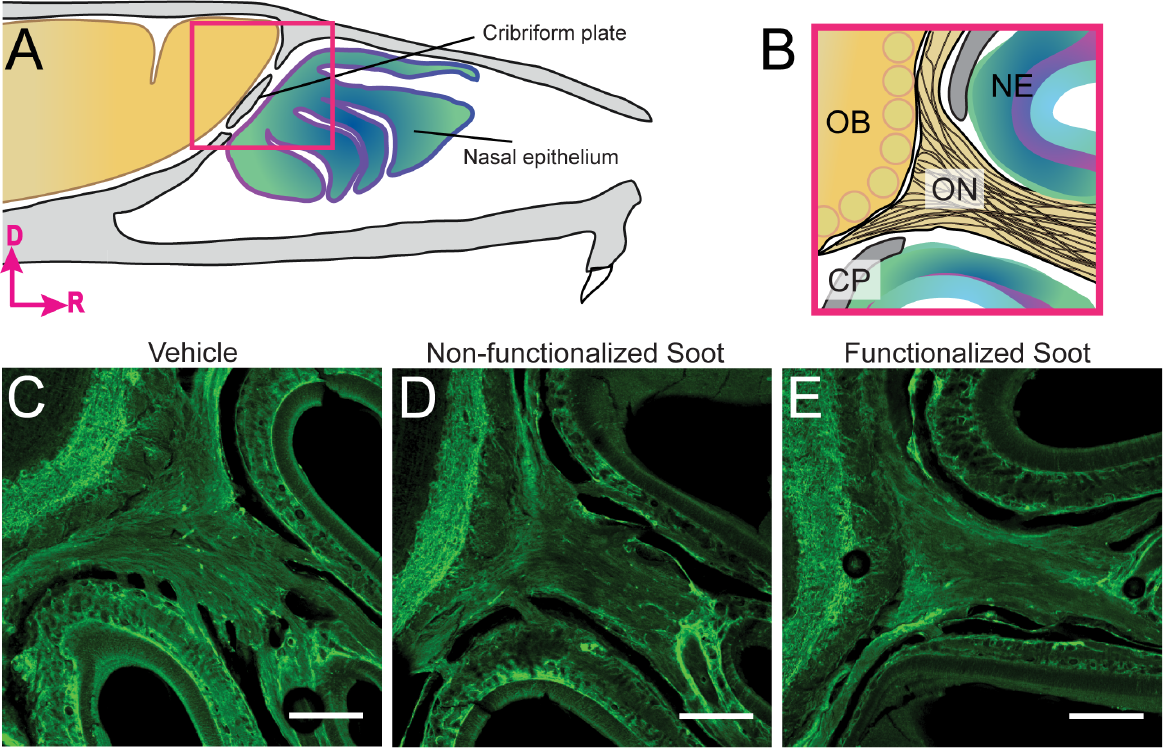
Soot treatment does not affect olfactory sensory neurons. A) Schematic showing sagittal cross-section of the mouse skull. Olfactory nerve (ON), neuroepithelium (NE), cribriform plate (CP), olfactory bulb (OB). B) Enlargement of box in A), showing the olfactory bulb, cribriform plate and foramina where the olfactory nerve enters the brain from the nasal cavity. Glomeruli are yellow circles. Example images of sagittal sections of the olfactory nerve location shown in B) from mice treated with vehicle (C), non-functionalized soot (D), and functionalized soot (E). Scale bar 250 µm.

## Methods

### Synthesis of soots

Synthetic soot was produced by functionalizing commercial carbon black, (Regal 250, Cabot Corp.). Carbon black was selected for its chemical purity and size similarity to diesel engine-produced soot. To introduce oxygen functional groups such as phenol, carboxyl and carboxylic, we used wet-chemical treatment based on acid etching (Romanos et al., 2011). In this preparation a gram of carbon black was treated with 100 mL laboratory grade concentrated nitric acid (HNO_3_, > 90%) under reflux for a duration of 24 hours at 80°C, just below the acid’s boiling point of 83°C. The carbon-acid mixture was continuously stirred using a magnetic stirrer to ensure uniform exposure and functionalization. The mixture was maintained at a consistent simmer and was thereafter washed with distilled water, filtered, and dried to obtain functionalized carbon black as a synthetic, oxidized soot. Any potential residual organic or aromatic compound present on the manufactured material as supplied would be oxidized and removed under these conditions.

### Characterization of soots

To visualize soot particles, we used transmission electron microscopy (FEI Talos F200X instrument equipped with Quad element EDS detector capable of both transmission electron microscopy (TEM) and scanning transmission electron microscopy (STEM)). For imaging, a beam acceleration voltage of 200 keV was used. Beam current was kept less than 5 nA for which sample damage or alteration is negligible at these magnifications (< 120 kX). Image defocus was one or two steps before the eucentric position. Images were captured using a Ceta-cooled CCD. Samples were dispersed and sonicated in methanol before being dropped onto 300 mesh C/Cu lacey TEM grids. High angle dark field (HAADF) images were obtained using an annular detector. EDS for elemental analysis and mapping was performed in the TEM. We also used the STEM mode, which has a high spatial resolution on the order of the minimum probe size (1.6 Å). The instrument was fitted with a 4-quadrant SDD Super-X EDS detector for EDS. The detection limit is typically < 1 atomic percent (at. %) depending on collection parameters. Typically, 7–10 regions of each material (nascent and functionalized forms) were sampled to gauge elemental representation. EDS was performed in STEM mode with a sample holder designed to provide low background signal for EDS. XPS experiments were performed using a Physical Electronics VersaProbe II instrument equipped with a monochromatic Al K_α_ x-ray source (hν = 1,486.7 eV) and a concentric hemispherical analyzer. Charge neutralization was performed using both low energy electrons (< 5 eV) and argon ions. Peaks were charge referenced to C-C band in the carbon 1s spectra at 284.5 eV. Measurements were made at a takeoff angle of 45° with respect to the sample surface plane. This resulted in a typical sampling depth of 3-6 nm (95% of the signal originated from this depth or shallower). Quantification was done using instrumental relative sensitivity factors (RSFs) that account for the X-ray cross section and inelastic mean free path of the electrons. A Thermogravimetric Analyzer (TA 5500, TA instruments) coupled to a Discovery Mass Spectrometer (MS) was used to analyze mass loss and the composition of the evolved gases as a function of temperature. The temperature was ramped up at 5°C/min in an inert atmosphere. The TGA features low volume, maximum temperature to 1200°C and has an inert quartz liner. The MS is a quadrupole mass spectrometer with a heated capillary interface, offering a 1-300 AMU range, unit m/z resolution. A Horiba LabRam Raman microscope was used to obtain Raman spectra for the samples when exposed to a 488 nm 100 mW laser with a 300 grooves/mm grating, providing a spectral resolution of 4 cm^-1^.

XPS was applied to dispersed powder to quantify both surface oxygen atom content (at. % basis) and distribution of oxygen functional groups (-C-OH, phenolic, -C=O, carbonyl, and -COOH, carboxylic), the nominal C1s (energy loss) positions were 286, 287 and 288.5 keV. CASA was applied to deconvolve the high-resolution spectra, with group contributions ratioed to the total oxygen elemental content. As a baseline, nascent (untreated) carbon black was also subject to the same analytical procedure as a “blank” sample. Wet acid reflux treatment of carbon black yielded ∼31 atomic % (near-surface) oxygen compared to the untreated carbon black, registering negligible surface content, (<1 at. %). By curve-fitting the C1s spectral loss profile, the calculated distribution across function groups was determined as 10.2% (phenolic, C-OH); 4.9% (carbonyl, C=O); and 9.4% (carboxylic, -COOH) (Vander Wal et al., 2011). (The good agreement (± 10%) in the measured and calculated value of atomic oxygen indicates appropriate curve fitting for functional group identification.) The TGA curve shows distinct regions of mass loss owing to functional groups leaving as temperature increases. The 5 wt.% net mass loss corresponds to the gasification of the carbon by the chemisorbed oxygen groups. Resolved by temperature, the TGA spectrum supported XPS identification of functional groups by successive mass loss stages for the oxygen group classes. Temperature resolved mass loss curves reveal m/z peaks at 44 AMU (CO_2_) arising predominantly from carboxylic groups and at 28 AMU (CO) arising from carbonyl and phenol groups (Kundu et al., 2008).

### Soot treatment protocol for mice

After the mouse had been rendered unconscious by a brief exposure to isoflurane, 20 µL of soot (functionalized or non-functionalized, 1% in sterile H_2_O) or vehicle control (sterile H_2_O) was administered to the left nare dropwise using a pipette. The animal was then inverted to allow for excess fluid to exit the nasal cavity. This treatment was repeated every other day (3 days a week) for one month. The animals were monitored and weighed daily after treatment.

### Histology

Mice were sacrificed via isoflurane overdose and perfused intracardially with heparinized-saline followed by 4% paraformaldehyde. The heads were fixed in 4% paraformaldehyde for 24 hours, decalcified for 48 hours in formic acid (4M) solution, and saturated in 30% sucrose for sectioning. Tissue sections of 100 µm thickness were sectioned on a freezing microtome. All staining was done in 24 well plates, with one section per well. Primary antibodies (and their respective dilutions) used on tissue sections were as follows: OMP (WAKO Chemicals U.S.A, 1:500), Pax6 (Santa Cruz, 1:250), and p63 (Abcam, 1:500). Sections were incubated in the primary antibody solution (primary antibodies + PBS-Triton) overnight at 4°C. Sections were then incubated in secondary antibody solution (secondary antibodies + PBS-Triton) for one hour at room temperature. The following secondary antibodies were purchased from Abcam and used at a 1:500 working dilution: Goat Anti-Rabbit IgG H and L (Alexa Fluor 488), Donkey Anti-Goat IgG H and L (Alexa Fluor 488), Goat Anti-Mouse IgG H and L (Alexa Fluor 488), Donkey Anti-Rabbit IgG H and L (Alexa Fluor 647), Donkey Anti-Goat IgG H and L (Alexa Fluor 647), and Goat Anti-Mouse IgG H and L (Alexa Fluor 647) preabsorbed. Sections were mounted on silane-coated unifrost slides (Azer Scientific), then cover slipped using fluoroshield mounting medium with DAPI (Abcam). Imaging was done on an Olympus Fluoview 1000 confocal and images were processed using ImageJ (NIH).

### Cell quantification procedures

To quantify the mean fluorescence of Pax6 and p63 antibody expression, images were first obtained on the Olympus FluoView 1000 confocal. Imaging settings were kept constant across samples to enable quantification of fluorescence. Using ImageJ (NIH), a rectangular ROI was drawn (250 µm in width and 50 µm in height) along the apical side of the neuroepithelium. For every animal, the ROI was drawn 250 µm in the rostral direction from the cribriform plate within the neuroepithelium located on the dorsal side of the medial olfactory nerve. The mean fluorescence of the ROI for each color channel (corresponding to each of the antibodies used) was obtained and averaged together for each treatment group. Data was plotted and analyzed in GraphPad Prism 8, using one-way ANOVA to test for significance.

### Quantification of mouse locomotion and rearing behavior

To measure any effects of intranasal soot treatment on behavior, mice were individually placed in a 34 × 31 × 14 cm (L x W x H) plastic box one month after the start of the treatment. All experiments were performed between 900 and 1600 ZT. The acquisition and analysis were done with the experimenter blinded to the treatment, and the order of animals was randomized. Mice were placed in the enclosure for 20 minutes, and the behavior was quantified over this entire period. The enclosure was cleaned with 70% ethanol between mice. The amount of locomotion and rearing behavior were monitored using an Intel® RealSense™ Depth Camera D435 (Hong et al., 2015). This camera provides simultaneous visible light and depth information used to calculate the animal’s distance from the camera. Images were acquired at a nominal rate of 15 frames/second using MATLAB (https://github.com/IntelRealSense/librealsense). To track the distance the animal traveled, the distance between the centroid of the mouse was calculated between each successive frame. This distance between frames is then summed over the course of the 20 minutes. Rearing events were defined as when the mean of the highest 20% of pixels of the mouse exceeded 8 cm from the bottom of the enclosure.

A generalized linear mixed-effects model (MATLAB function *fitglme*) was used to evaluate the differences in rearing events, rearing duration, and distance traveled. Each treatment (vehicle, non-functionalized and functionalized soot) was a fixed-effect, with the sex treated as a random effect.

### Data availability

Code for the acquisition, analysis and plotting of the behavioral data, as is the behavioral data plotted in figure 4, is available here: https://github.com/DrewLab/Norwood_Gharpure_Turner_Ferrer-Pistone_VanderWal_Drew_Manuscript2020

**Figure 4:**
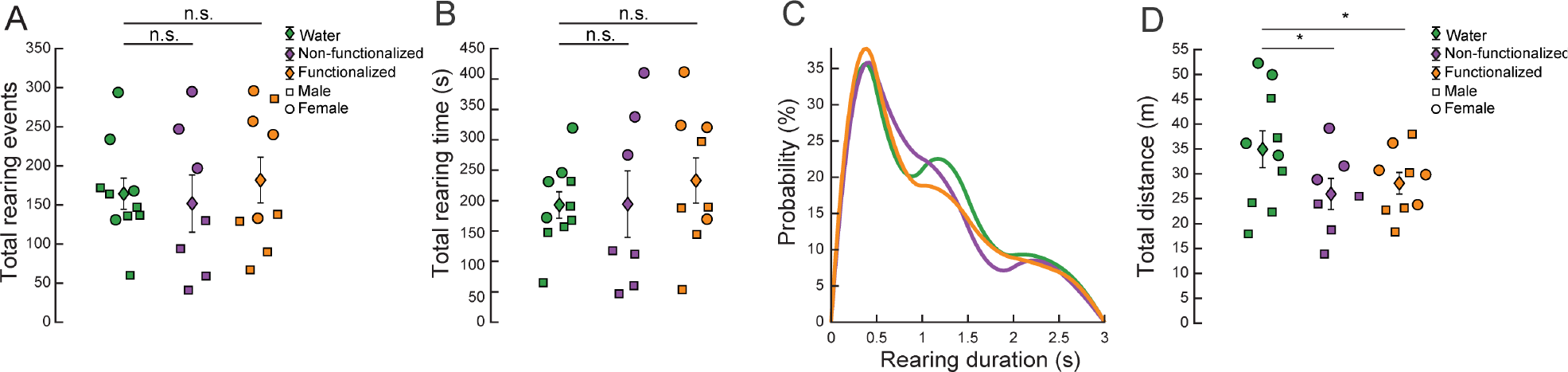
Soot treatment has minimal effects on locomotion and rearing behavior. Comparisons of the effects of vehicle, non-functionalized, and functionalized soot on spontaneous rearing and locomotion behaviors over 20 minutes. The data from each individual mouse is shown as a square (males) or circle (females). The mean of each group is shown as a diamond, standard deviation is denoted with error bars. A) Plot of total number of rearing events for each treatment type. There was no significant difference in the number of rearing events between the control and either of the soot treatment groups (p < 0.62 non-functionalized, p < 0.63 functionalized). B) Plot of total rearing time. There was no significant difference in the total rearing time between groups (p < 0.94 non-functionalized, p < 0.33 functionalized). C) Probability distribution of individual rearing event durations for each of the treatment groups. (D) Plot of total distance travelled by each mouse. Treatment with non-functionalized soot (p < 0.02) and functionalized soot (p < 0.05) both significantly decreased the total distance traveled relative to the vehicle treated group.

## Results

### Characterization of soot compounds

Transmission electron micrograph of a carbon black aggregate and primary particle is shown in Fig. 1. The aggregate consists of pseudo-spherical primary particles, partially merged or fused together forming a fractal aggregate. A Raman spectrum of the nascent carbon black is shown in Fig. 1c. Raman spectroscopy has been developed as a standard method for determining the planar coherence lengths (L_a_) in graphitic carbon, which possesses limited long-range order (Tuinstra and Koenig, 1970). The lower frequency “D” peak at ∼1360 cm^-1^ arises from disorder-induced Raman activity of zone-boundary A_1g_ phonons whereas the “G” peak at ∼1580 cm^-1^ reflects the in-plane stretching motion of the aromatic rings, designated as E_2g_ motions. Their comparable intensity reveals considerable disordered carbon, characteristic of furnace blacks and representative of combustion-produced soot emissions (Dennison et al., 1996; Sadezky et al., 2005). Their intensity ratio is an accepted technique for determining L_a_ in disordered graphitic materials, given by the relation 4.4(I_d_/I_g_)^-1^ = L_a_, calculated here as 0.85 nm, a value commensurate with the short lamellae viewed by HRTEM (Sadezky et al., 2005). The asymmetry of the D-peak due to the extended low frequency (shift) tail is consistent with further disorder of the carbon lattice such as sp^3^ and sp^2^ carbon at the periphery of the crystallites, contributing vibrations of A_1g_ symmetry (Sadezky et al., 2005; Parent et al., 2016). Fluorescence from the oxygen groups and their auxochromic interactions with the *π* electrons of the sp^2^ carbon network dwarfed the Raman signature of the functionalized material, preventing its comparison to the nascent material.

Figure 1D shows a high angle dark field TEM image of soot particles for reference and respective EDS map displaying carbon (blue) and oxygen (red) for the nitric acid functionalized carbon black. While EDS cannot point to a definitive volumetric vs. surface oxygen presence given its 2-D nature, nitric acid–treated carbon black shows oxygen appearing to be concentrated along the particle perimeter, reflecting a higher near-surface contribution near the particle edge along the beam path. In the 2-D image it must be noted that EDS shows relative amounts of elemental carbon and oxygen and does not give information on what functional groups are present. An image of the 1% solution of soot in water that is applied intranasally is shown in Figure 1E.

### Soot accumulates in the nasal passageway and lungs, but does not change the structure of the olfactory nerve or OSNs

We treated mice intranasally with vehicle, non-functionalized soot, or functionalized soot for one month. Mice were briefly anaesthetized with isoflurane and an intranasal solution of soot (functionalized or non-functionalized, 1% in sterile water) or vehicle (sterile H2O) was administered to the left nare. This treatment was repeated every other day (three times a week) for one month. We saw no appreciable differences between weight of mice of the different treatment groups (data not shown). After sacrifice, the skulls were rapidly decalcified (Norwood et al., 2019) and sectioned. Examples of thin sections of olfactory bulb/nasal cavity area are shown in Fig 2A-C, and accumulation of functionalized soot in the olfactory epithelium, but not non-functionalized soot was observed. Soot could be seen in the lungs of treated animals (Fig 2D-F). A total of 0.2mg of soot particles was applied each day, though we conservatively estimate that < 20% remained in the nose after inverting the mouse. Given that an average mouse respiratory volume over a day is ∼34.5 L (0.15 mL tidal volume with 160 breaths a minute (Zhang et al., 2019)) this works out an effective dose equivalent to breathing air with a PM_2.5_ level of ∼300 µg/m^3^, comparable to the air quality in Beijing (Zíková et al., 2016) or New Delhi (Pant et al., 2017).

We also examined the status of olfactory sensory neurons to see if either type of soot had a detectable effect on their health. To assess any damage to the olfactory bulb or nerve caused by exposure to the soot particles, the area was examined histologically utilizing rapid decalcification and sectioning. Olfactory sensory nerves enter the cranial compartment through the cribriform plate (Bird et al., 2018). We visualized the nerve in soot and vehicle-treated mice using an antibody against olfactory marker protein (OMP) (Fig. 3). We found no discernable difference in olfactory nerve labeling among the treatments, indicating that soot treatment does not have any obvious effect on olfactory sensory nerves for the treatment duration used. This is markedly different from intranasal treatment with zinc sulfate, a single treatment which causes the rapid and irreversible ablation of olfactory sensory neurons (Norwood et al., 2019).

### Effects of intranasal soot treatment on exploratory behavior

One important question is to what extent soot treatment impacts the behavior of the mice. We quantified locomotion and rearing behavior using an Intel RealSense D435 depth-sensing camera (Hong et al., 2015) after intranasal soot exposure (Fig. 4). Treated mice from all three treatment groups were individually placed in a novel environment (white plastic container) after the one month of treatment and their movement and rearing behaviors were monitored for 20 minutes. Rearing behavior is a measure of anxiety (Sturman et al., 2018), and locomotion can be used to assay sickness and malaise (Engeland et al., 2001). No significant differences were observed in total rearing events or total rearing time for all treatment groups. However, a significant difference in total distance traveled was observed between the vehicle and both the functionalized and non-functionalized soot treatment groups. If the soot treatment causes pronounced health problems, we might expect large decreases in the amount of time rearing or locomotion behavior. As we did not observe pronounced changes in behaviors, this suggest the soot treatment does not cause any generalized decreases in health.

### Functionalized soot decreases the number of olfactory progenitor cells

As we saw no obvious changes in olfactory sensory neurons and their axons, we then asked how soot treatment might affect other cell types in the nasal epithelium, particularly the progenitor cells that directly and indirectly give rise to olfactory sensory neurons. If these cells are damaged, then this could lead to a long-term decline in the number of OSNs as the animals age. We used immunofluorescence staining of the neuroepithelium to visualize changes in progenitor cells (Fig. 5A-B). The expression of the anti-Pax6 or anti-p63 primary antibody in the neuroepithelium was quantified to assess any disruptions in the number of globular basal cells (GBCs) or horizontal basal cells (HBCs), respectively (Fig. 5C-G). In the neuroepithelium, compared to the non-functionalized and vehicle treatments, a significant decrease in the number of Pax6^+^ cells was observed following the one-month treatment with functionalized soot (Fig. 5C-F). This decrease in Pax6^+^ cells implies a decrease in the pool of OSN progenitor cells. However, no significant differences in the number of p63^+^ cells were observed in the neuroepithelium for any of the treatment types (Fig. 5C-E, G), indicating no response by the HBCs following soot exposure. Decreases in the number of GBCs could lead to decreases in the number of olfactory sensory neurons in the long term.

**Figure 5:**
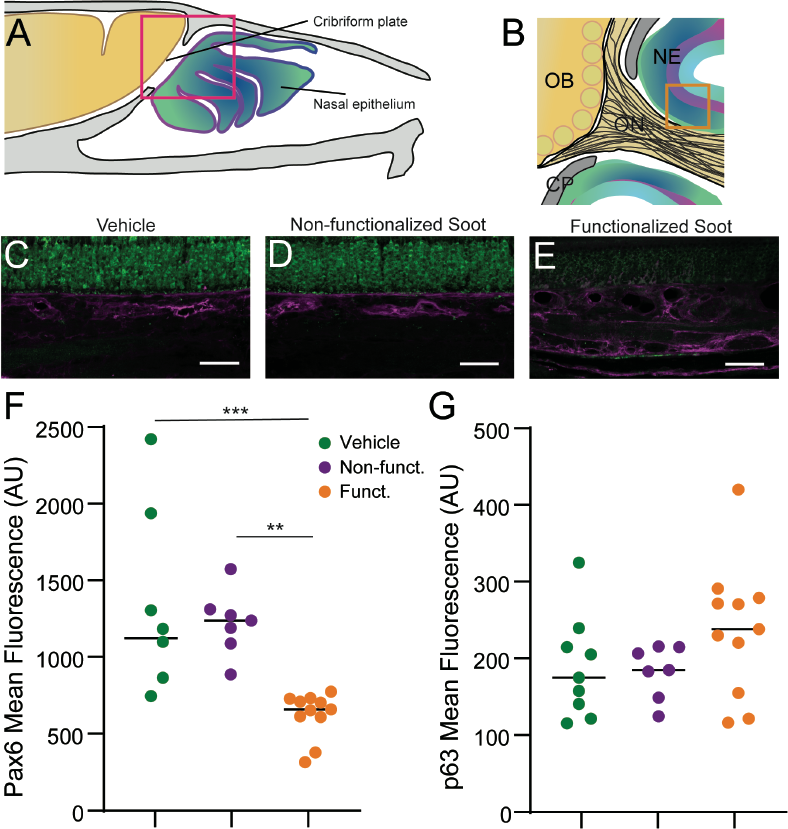
Functionalized soot treatment decreases the number of Pax6^+^ GBC olfactory progenitor cells. A) Schematic of the sagittal plane of the mouse cranial and nasal cavities. Olfactory nerve (ON), neuroepithelium (NE), cribriform plate (CP), and olfactory bulb (OB). Glomeruli are yellow circles. B) Sagittal view of the area within the pink box in A. C-E) Immunofluorescent staining of an area enclosed by the orange box. Pax6 is shown in green and p63 is shown in magenta. Scale bar 250µm. C) Vehicle control. D) Non-functionalized soot. E) Functionalized soot. F) Quantification Pax6 fluorescence. Functionalized soot caused a decrease in Pax6 expression (one way ANOVA, F(2,24) = 11.04, p < 0.0004) relative to non-functionalized soot (post-hoc unpaired t-test, t(24) = 3.572, p < 0.0046) and vehicle control (post-hoc unpaired t-test, t(24) = 4.266, p < 0.00008). G) No significant difference between group means of p63 fluorescence (one-way ANOVA, F(2,24) = 1.797, p < 0.19).

## Discussion

In order to understand how air pollution might affect olfactory sensory neurons and their progenitor cells, we treated mice intranasally with surrogate soot-like particles that either had oxygen-functionalized surfaces or non-functionalized surfaces. We found that these compounds had minimal effects on behavior, the olfactory sensory nerve, or horizontal basal cells. However, oxygen functionalized soot greatly reduced the population of globular basal cells. Our results are consistent with many other studies that have found that oxidized soots are more cytotoxic than un-oxidized soots (Li et al., 2009; Holder et al., 2012; Pourkhesalian et al., 2015).

Our results suggest a potential model of how long-term exposure to air pollutants could drive anosmia and decreased CSF outflow into the nasal cavity (Fig. 6). Exposure to oxidized soot particles reduces the number of GBCs. As olfactory sensory neurons senesce, the reduced population of GBCs leads to incomplete replacement of OSNs. The decrease in OSNs could then potentially lead to decreases in olfactory sensitivity seen with exposure to air pollution (Ajmani et al., 2016a; Ajmani et al., 2016b; Hummel et al., 2017). The decrease in OSN axons could also reduce fluid outflow through the cribriform plate (Norwood et al., 2019), which may lead to slower CSF turnover and a buildup of toxic metabolites in the CSF, and potentially to development of neurodegenerative diseases (Iliff et al., 2012). We emphasize though this explanation unites many disparate observations, it is nonetheless speculative.

**Figure 6:**
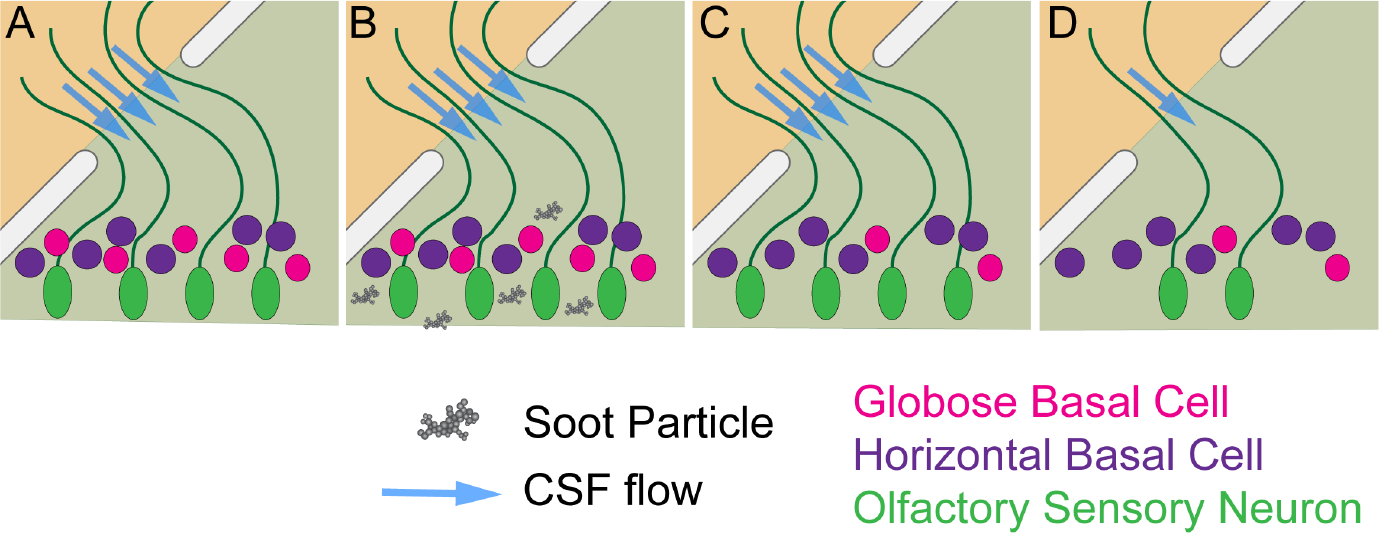
Potential impact of decreased number of GBCs on CSF outflow. A) Normally, cerebrospinal fluid drains (between olfactory sensory neuron axons) out of the cranial compartment (upper left) into the nasal compartment. Exposure to oxidized soots (B) results in the death of GBCs (C). Decreases in the number of GBCs eventually leads to a decrease in the number of olfactory sensory neurons (D), as they are not replaced after they die, so there is less CSF draining out of the cribriform plate.

There are several limitations to our study. We do not know the mechanism by which oxygen functionalized soot preferentially damages GBCs. It could be that oxygen functionalized soot is more prone to accumulating in the nasal epithelium (Fig. 2), enhancing their toxic effects. Only a single kind of air pollutant was tested here, and other known pollutants (such as heavy metals) could have different or synergistic effects (Calderón-Garcidueñas et al., 2002; Costa et al., 2020). Finally, we only looked at relatively acute effects of exposure, and longer-term exposure may have different effects.

## Acknowledgements

This work supported by an PSU Institute of Energy and Environment seed grant to RVW and PJD, F31NS105461 to JNN, and R01NS078168 to PJD.

## Competing Interests

None

## Author contributions

JNN, APG, KLT and LFP performed experiments, JNN, APG and KLT analyzed data, JNN, RVW and PJD designed the experiments and wrote the manuscript with input from all authors.

## Notes

### Competing Interest Statement

The authors have declared no competing interest.

